# Altered Cardiolipin Metabolism is Associated with Cardiac Mitochondrial Dysfunction in Pulmonary Vascular Remodeled Perinatal Rat Pups

**DOI:** 10.1101/2021.10.12.464115

**Authors:** Laura K. Cole, Genevieve C. Sparagna, Vernon W. Dolinsky, Grant M. Hatch

**Affiliations:** Department of Pharmacology and Therapeutics, Children’s Hospital Research Institute of Manitoba, Rady Faculty of Health Sciences, University of Manitoba, Winnipeg, Canada; Diabetes Research Envisioned and Accomplished in Manitoba (DREAM) Theme, Children’s Hospital Research Institute of Manitoba, Rady Faculty of Health Sciences, University of Manitoba, Winnipeg, Canada; Department of Medicine, Division of Cardiology, University of Colorado Anschutz Medical Center, Aurora, Colorado, USA

## Abstract

**Background:** Pulmonary vascular remodeling (PVR) *in utero* results in the development of heart failure (HF). The alterations that occur in cardiac lipid and mitochondrial bioenergetics during the development of *in utero* PVR was unknown.

**Methods:** PVR was induced in pups *in utero* by exposure of pregnant dams to indomethacin and hypoxia. Cardiac lipids, echocardiographic function and cardiomyocyte mitochondrial function were subsequently examined.

**Results:** Perinatal rat pups with PVR exhibited elevated left and right cardiac ventricular internal dimensions and reduced ejection fraction and fractional shortening compared to controls. Cardiac myocytes from these pups exhibited increased glycolytic capacity and glycolytic reserve compared to controls. However, respiration with glucose as substrate was unaltered. Fatty acid oxidation and ATP-insensitive respiration were increased in isolated cardiac myocytes from these pups compared to controls indicating mitochondrial dysfunction. Although abundance of mitochondrial respiratory complexes were unaltered, increased trilinoleoyl-lysocardiolipin levels in these pups was observed. A compensatory increase in both cardiolipin (CL) and phosphatidylethanolamine (PE) content were observed due to increased synthesis of these phospholipids.

**Conclusion:** Alterations in cardiac cardiolipin and phospholipid metabolism in PVR rat pups is associated with the mitochondrial bioenergetic and cardiac functional defects observed in their hearts.

**Impact statement:**

- Phospholipid metabolism was examined in pulmonary vascular remodeling in perinatal rat pups.
- Pulmonary vascular remodeling was induced *in utero* by treating pregnant dams with hypoxia and indomethacin at 19-21 days of gestation.
- The offspring exhibited altered pulmonary arterial remodeling with subsequent cardiac hypertrophy, ventricular dysfunction, cardiac myocyte mitochondrial dysfunction with altered fatty acid utilization.
- In addition, the offspring exhibited elevated cardiolipin, lysocardiolipin and phosphatidylethanolamine content which may potentially contribute to the cardiac mitochondrial dysfunction.

## Introduction

Pulmonary hypertension of the newborn is a failure of normal pulmonary vascular relaxation after birth, with incidence of up to 6 per 1000 live births and 10-30% mortality (1–3). It is a significant cause of heart failure (HF) in healthy newborn infants. In healthy term infants it is caused by perinatal hypoxia, inflammation or direct lung injury. These infants develop the HF rapidly within a week and HF becomes the main limiting factor for their survival. A number of studies in human and animal models have demonstrated alterations in mitochondrial bioenergetics during the development of HF (4). Several of these studies have implicated alterations in the mitochondrial phospholipid cardiolipin (CL) as a contributor to the mitochondrial dysfunction. CL, the signature phospholipid of mitochondria, is essential for mitochondrial morphology, bioenergetics, dynamics, and signaling pathways (5–11). However, limited information exists on the changes in cardiac function and phospholipid composition that occur during *in utero* pulmonary vascular remodeling (PVR). Previously we demonstrated that newborn piglets exposed to a hypoxic environment for 3 days developed alterations in CL and reduced mitochondrial respiratory chain dysfunction during the development of a right ventricular hypertrophy (12).

In this study, we utilized a unique gestational rat model PVR to examine how cardiac phospholipid metabolism, cardiac morphology and mitochondrial function are impacted by PVR. We show for the first time that CL and phosphatidylethanolamine (PE) levels are elevated in the hearts of PVR rat pups and that this is accompanied by an accumulation of lysocardiolipin (LysoCL) and mitochondrial bioenergetic and cardiac dysfunction.

## Materials and Methods

### Animals

This study was performed with approval of the University of Manitoba Animal Policy and Welfare Committee which adheres to the principles for biomedical research involving animals developed by the Canadian Council on Animal Care and the Council for International Organizations of Medical Sciences. This study is reported in accordance with ARRIVE guidelines. All animals were maintained in an environmentally controlled facility (22°C, 37% humidity, 12 h light/dark cycle) with free access to food and water.

PVR was induced in perinatal rats by treating pregnant dams during 19-21 days of gestation with hypoxia and indomethacin (single ip dose 0.5 mg/Kg in sterile PBS pH 7.4) as described (13, 14). Timed pregnant rats at 18 days gestation were utilized. One set of pregnant rats were housed normally (Control, single ip dose sterile PBS) as control animals, a second set of pregnant rats were indomethacin-treated as above (Indo), while a third set of pregnant rats were indomethacin-treated and then housed in a hypoxic environment (12% oxygen) (PVR). This was accomplished by housing rats in a plexiglass chamber and the chamber was maintained at 12% oxygen using a nitrogen washout system. Medical grade compressed oxygen (12% oxygen balance Nitrogen, Welder’s Supply) was used in combination with medical grade nitrogen to adjust chamber oxygen levels to 12% using a gas analyzer (Radiometer ABL 700 series). Compressed gas flowed continuously into the chamber at a rate of 1 L/min for 3 days. Chamber oxygen levels, flow rates and temperatures were monitored every 24 h at a minimum. On the fourth day all animals were sacrificed and caesarean section was performed to remove the pups. Prior to sacrifice fetal echocardiographic parameters were determined as described below. After removal by caesarean section the pups were then weighed, measured for length and the heart, liver and lung removed and weighed. Histology of pulmonary arteries was performed as previously described (12). In some experiments, the heart was freeze dried and weighed. In other experiments, freshly isolated hearts were used for preparation of mitochondrial fractions or isolation of cardiac myocytes as outlined below.

### *In vivo* echocardiography

Transthoracic echocardiography was performed on dams mildly anesthetized with 1-1.5% isoflurane, 1L/min oxygen as previously described (15). Each rat was placed on a heated ECG platform to maintain body temperature and obtain fetal heart measurements. A Vevo 2100 High resolution imaging system equipped with a 30-MHz transducer (Visual Sonics, Toronto) was used to visualize the fetal hearts. Measurements obtained included interventricular septum diastolic (IVSd), interventricular septum systolic (IVSs), left ventricle internal dimension diastolic (LVIDd), left ventricle internal dimension systolic (LVIDs), left ventricular wall diastolic (LVWd), intraventricular septum end systole (LVSs), ejection fraction (EF), fractional shortening (FS), right ventricular wall diastolic (RVWd), right ventricular wall systolic (RVWs), right ventricle internal dimension diastolic (RVIDd) and right ventricle internal dimension systolic (RVIDs).

### Cardiac myocyte preparation and radiolabeling experiments

Cardiac myocytes were prepared from the hearts of control and PVR pups as described (16). The protocol allows for rat cardiomyocytes to be cultured for up to 72 h at 37°C in 5% CO_2_ without significant change in phenotype (17). Enough hearts were collected to yield approximately 15 – 20 million cardiac myocytes sufficient to plate 15-20 × 35mm dishes (1 million/dish, Corning Primeria^TM^). Isolated cardiac myocytes were incubated with 0.1 mM [1,3-^3^H]glycerol (2 μCi/dish, Perkin Elmer) or 0.1 μM [1-^14^C]linoleic acid (2 μCi/dish, Perkin Elmer) bound to albumin (1:1 molar ratio) or 0.1 μM [1-^14^C]oleic acid (2 μCi/dish Perkin Elmer) bound to albumin (1:1 molar ratio) for up to 360 min (6 h) and radioactivity incorporated into phospholipids determined as previously described (18).

### Respiratory function analysis

The oxygen consumption rate (OCR) was measured from isolated cardiac myocytes of control and PVR pups (1 × 10^5^/well, coated with fibronectin) using a Seahorse XF24 Bioscience instrument (18). XF assay media contained either 1 mmol/L pyruvate and 25 mmol/L glucose for glucose metabolism or 1 mmol/L pyruvate, 2.5 mmol/L glucose, 0.5 μmol/L carnitine, and 0.175 mmol/L palmitate-BSA for fatty acid (FA) metabolism. Basal oxygen consumption was considered to be the basal respiration sensitive to inhibition by 1 μmol/L antimycin A plus 1 μmol/L rotenone. ATP-sensitive oxygen consumption was inhibited by 1 μmol/L oligomycin, and ATP-insensitive respiration (heat) was the remaining proportion of basal oxygen consumption. Maximal oxygen consumption was achieved with 1 μmol/L carbonyl cyanide 4-(trifluoromethoxy) phenylhydrazone. Fatty acid-dependent respiration was measured as the difference in oxygen consumption measured in the presence of 40 μmol/L etomoxir and vehicle (water). The XF glycolysis stress test kit (Aligent) was used for glycolysis analysis. Briefly, cells are cultured in the absence of glucose followed by the sequential addition of glucose (10mM), oligomycin (3μM) and 2-DG (1M). Where glycolysis is measured following glucose addition (glucose – 2-DG), glycolytic capacity measured following oligomycin addition (oligomycin – 2-DG), and glycolytic capacity the difference between them (glycolytic capacity-glucose).

### Western blot analysis of mitochondrial respiratory subunits

Cardiac mitochondria were isolated from control or PVR hearts using the MITOISO1 Mitochondria Isolation Kit (Sigma) as previously described (15). Mitochondrial protein (7.5 μg) from was separated on the Bio-Rad mini gel electophoresis system by SDS-PAGE (12% acrylamide) as previously described (15). Western blot antibody cocktail (Abcam) contained (55 kDa, anti-ATP synthase subunit ATP5a, Complex V; 47 kDa, anti-complex III subunit core 2 UQCRC2; 35 kDa, anti-complex IV subunit MTCO1; 30 kDa, complex II subunit SDHB; and 20 kDa, complex I subunit NDUFB8). Adult rat mitochondrial protein was added for comparative control and α-tubulin (Cell Signaling) was used as the loading control. Proteins were visualized by chemiluminescence using the ECL Western blotting detection system (Amersham).

### Lipid analysis

The level of the major phospholipids from cardiac tissue homogenates including phosphatidylcholine (PC), PE and CL were determined by HPLC separation followed by lipid phosphorus assay (19, 20). Molecular species of CLs and lysoCLs were quantitated from tissue homogenates by HPLC coupled to electrospray ionization mass spectrometry (21). Cardiac cholesterol, cholesterol ester and triacylglycerol were determined by HPLC as previously described (15).

### Statistical analysis

Data are expressed as means ± standard error of the mean (SEM). Comparisons between control and PVR offspring were determined using 1-way analysis of variance using Tukey post-hoc analysis. For each measurement, the offspring were derived from multiple litters. A probability p value of <0.05 was considered significant.

## Results

PVR was induced in perinatal rats by treating pregnant dams during 19-21 days of gestation with hypoxia and indomethacin (13, 14). Lung histology analysis of newborn rats revealed classic PVR in pups from the hypoxic-indomethacin treated group compared to control including increased medial thickness in the larger pulmonary arteries with no changes in their external diameter (Fig. 1A-D). Thus, we established this model of PVR in fetal rats (13, 14). Reduction in body weight and length of newborn PVR pups were accompanied by reductions in both lung and liver weight compared to control (Fig. 2A-D). Dried heart weight/body weight ratio was elevated in PVR pups compared to control (Fig. 2E) and this was accompanied alterations in cardiac structural and functional parameters (Fig. 3A-C). Specifically, in the left ventricle elevations in IVSd, IVSs, LVWd and LVSs were accompanied by a reduction in EF and FS. In addition, right ventricle dimensions including RVWd and RVWs were elevated in PVR pups compared to control. Thus, the PVR rat hearts exhibited cardiac hypertrophy and mechanical dysfunction.

**Figure 1.**
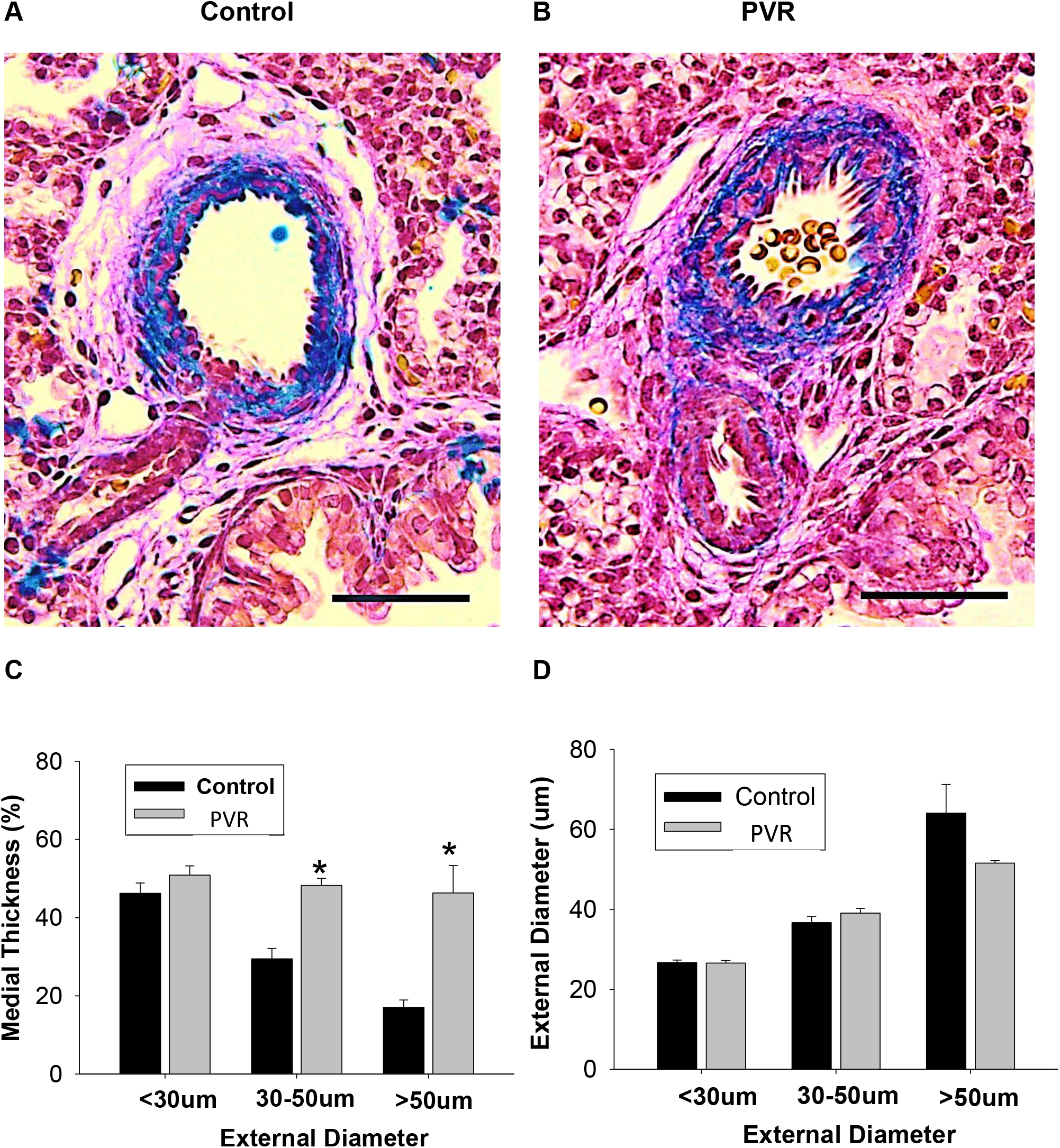
Histology of PVR arteries. Masson’s trichrome staining of control (A) and PVR (B) pulmonary arteries. Representative sections are depicted. C. Medial thickness, and D. External diameter of control and PVR pulmonary arteries. The black bar insert represents 50 μM. Data represent the mean ± SEM, n=5 Control, n=23 PVR, *p<0.05.

**Figure 2.**
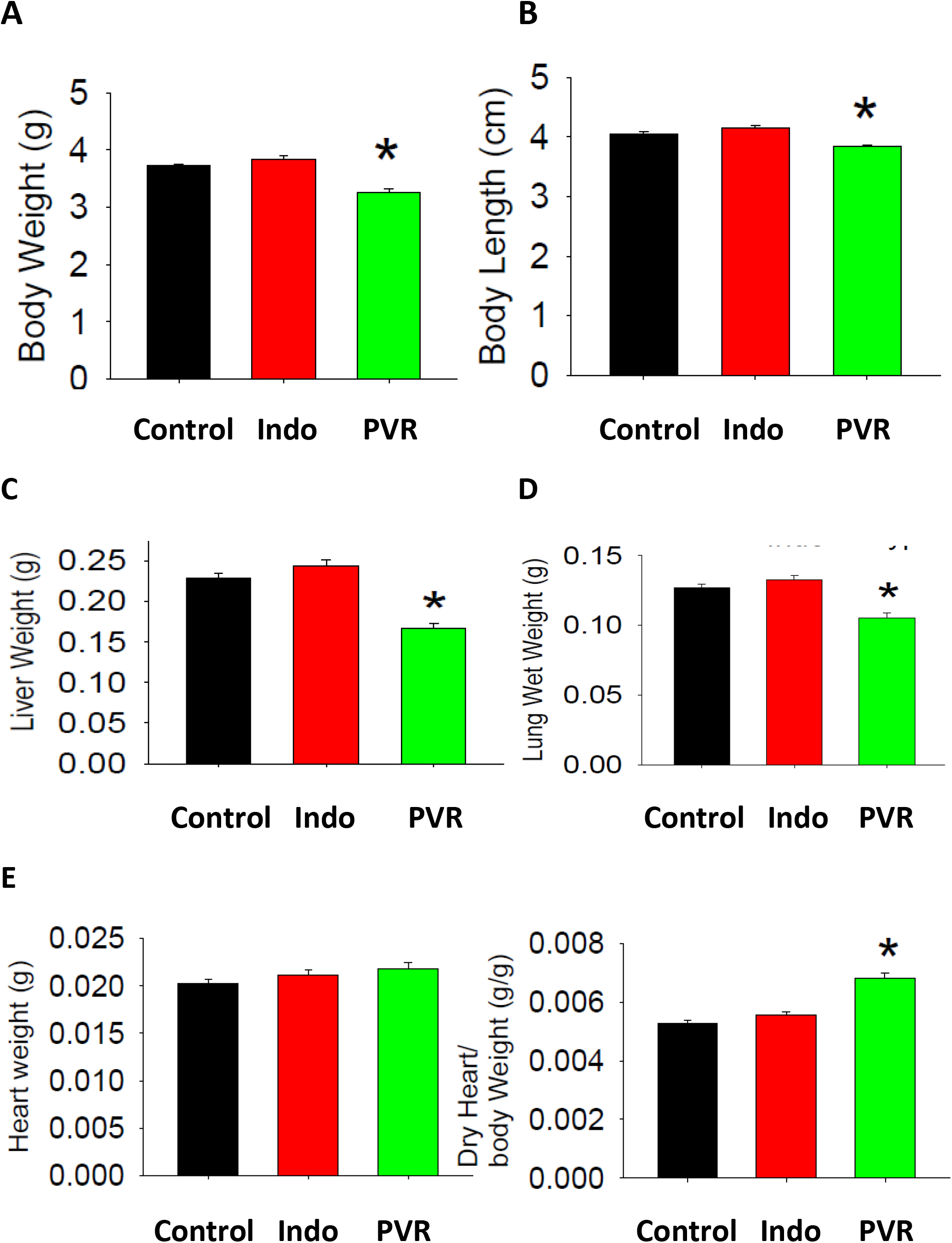
Body weight, length and organ weight of PVR rats. Whole body weight (A) and length (B) of control and PVR rat pups. C. Heart weight and dry heart weight/body weight ratio of control and PVR rats. Data represent the mean ± SEM, n=5 Control, n=23 PVR, *p<0.05.

**Figure 3.**
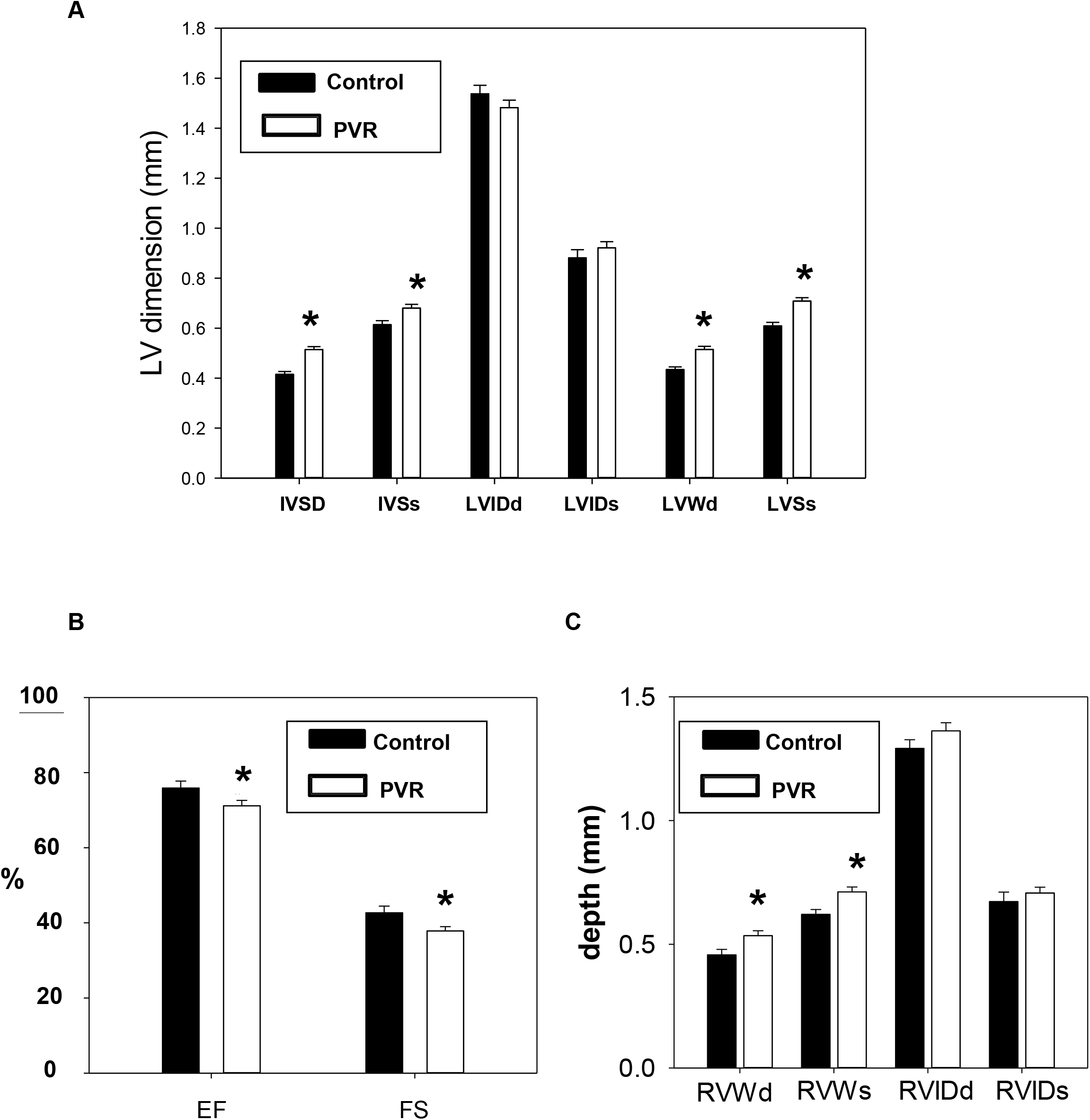
Echocardiographic parameters of PVR hearts. Transthoracic echocardiography was performed on control and PVR rats. A. Interventricular septum diastolic (IVSd), interventricular septum systolic (IVSs), left ventricle internal dimension diastolic (LVIDd), left ventricle internal dimension systolic (LVIDs), left ventricular wall diastolic (LVWd), intraventricular septum end systole (LVSs). B. Ejection fraction (EF) and fractional shortening (FS). C. Right ventricular wall diastolic (RVWd), right ventricular wall systolic (RVWs), right ventricle internal dimension diastolic (RVIDd) and right ventricle internal dimension systolic (RVIDs). Data represent the mean ± SEM, n=5 Control, n=23 PVR, *p<0.05.

We examined if substrate utilization was altered in isolated cardiac myocytes from rats in which PVR was induced during gestation. Oxygen consumption rate (OCR) was unaltered in PVR pups compared to control when glucose was used as substrate (Suppl. Fig. 1A). Interestingly, glycolytic capacity and glycolytic reserve were increased in PVR pups compared to control (Fig. 4A). In addition, when palmitate was used as substrate, an increase in basal OCR was observed which was blocked by incubation with the carnitine palmitoyltransferase-1 inhibitor etomoxir (Fig. 4B). The increase in fatty acid (FA) oxidation was accompanied an increase in ATP-sensitive oxygen consumption (Fig. 4C) and ATP-insensitive respiration (Fig. 4D) in PVR pups compared to control. Maximum OCR was unaltered in cardiac myocytes between control and PVR pups (Fig. 4E). Thus, the increase in ATP-insensitive respiration indicated a dysfunction in efficiency of mitochondrial FA oxidation in cardiac myocytes from PVR rat pups.

**Figure 4.**
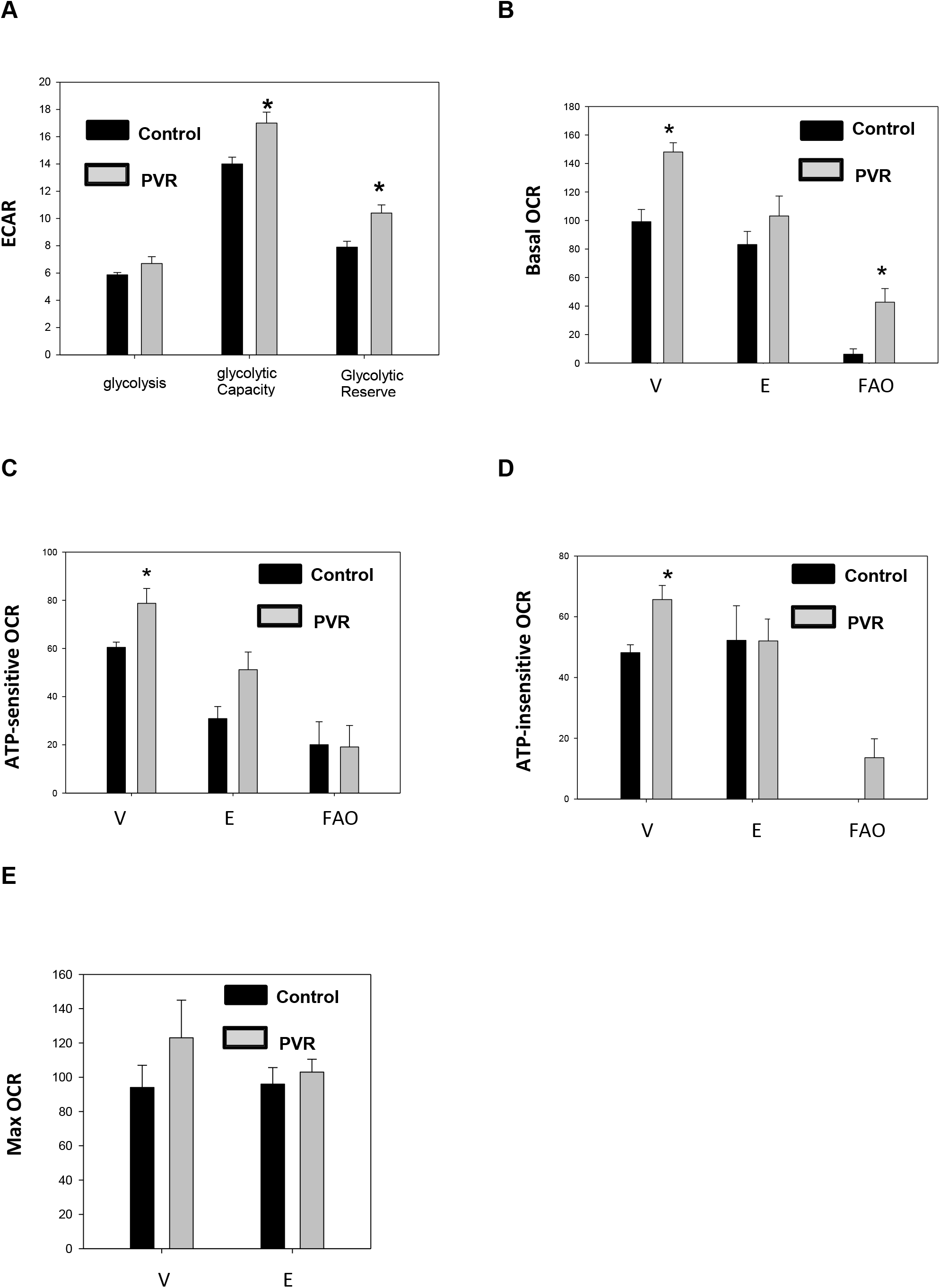
Mitochondrial dysfunction is observed in PVR rats. Glycolysis, glycolytic capacity and glycolytic reserve in control and PVR cardiomyocytes. ECAR, extracellular acidification rate (mPH/min/ug protein). Basal oxygen consumption rate (OCR) (pmol/min/ug protein) (B), ATP-sensitive OCR (pmol/min/ug protein) (C), ATP-insensitive OCR (pmol/min/ug protein) (D) and maximum (Max) OCR (pmol/min/ug protein) (E) in control and PVR cardiomyocytes. V, vehicle; E, plus etomoxir; FAO, calculated fatty acid oxidation. Data represent the mean ± SEM of n=3, *p<0.05.

We next examined the pool sizes of the three major phospholipids in hearts from control and PVR pups. Phospholipid analysis revealed that there was no alteration in the levels of phosphatidylcholine (PC) but striking increases in both cardiolipin (CL) and phosphatidylethanolamine (PE) in hearts of PVR pups compared to control (Fig. 5A-C). The observed reduction in cardiac PC/PE ratio was consistent with that seen in pressure-induced heart failure (22). In addition, the levels of cardiac cholesterol, cholesterol ester and triacylglycerol were unaltered (Suppl. Fig. 2A-C). Since CL, and specifically tetralinoleoylcardiolipin (L_4_CL), are required for optimal cardiac mitochondrial bioenergetic function (23), we examined the molecular species composition of CL. Elevations in 1442, 1448 and 1472 species of CL appeared to be responsible for the observed increase in CL in PVR pups (Fig. 5D). The highest being linoleate containing L_4_CL (1448). In isolated cardiac myocytes from PVR pups the increase in L_4_CL was due to an increase in [1-^14^C]linoleate incorporation into CL indicating increased synthesis from linoleate (Fig 6A). In contrast, *de novo* synthesis of CL from [1,3-^3^H]glycerol was unaltered in cardiac myocytes from PVR rat pups (Fig. 6B).

**Figure 5.**
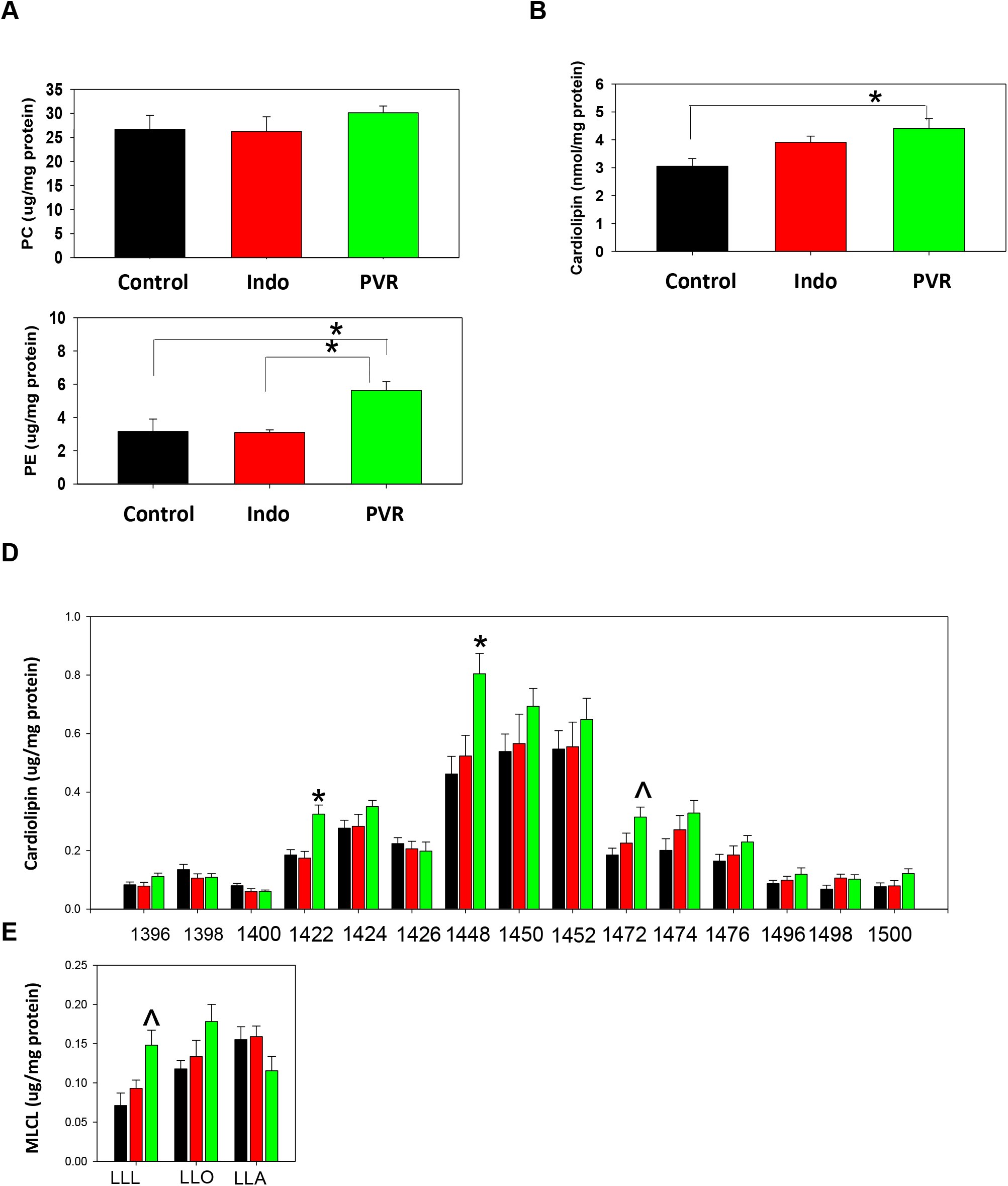
PE and CL are elevated in PVR rat hearts. The content of PC (A), CL (B) and PE (C) and CL molecular species composition (D) and lysoCL molecular species composition (E) were determined in hearts of newborns from control, indomethacin (Indo) or PVR rats. D. the numbers represent individual CL molecular species. E. LLL, L_3_-lysocardiolipin; LLO, linoleoyl, linoloeyl, oleoyl-lysoCL; LLA, linoleoyl, linoleoyl, arachidonyl-lysoCL. In D and E: Control, black bars; Indo, red bars; PVR, green bars. Data represent the mean ± SEM of n=3, *p<0.05.

**Figure 6.**
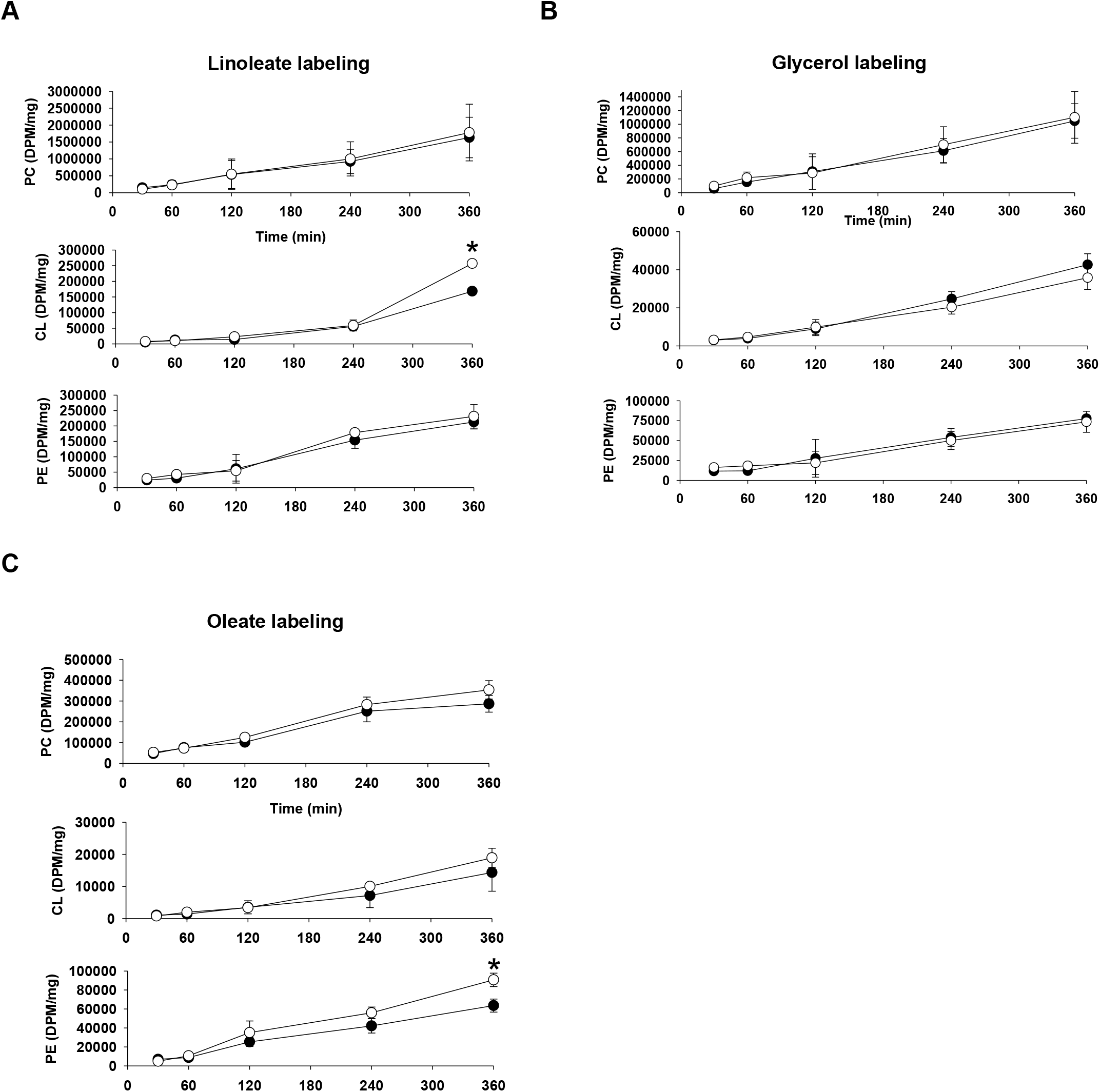
Synthesis of phospholipids from glycerol, linoleate and oleate in isolated PVR cardiac myocytes. Isolated control or PVR cardiomyocytes were incubated for up to 360 min with [1-^14^C]linoleate (A), [1,3-^3^H]glycerol (B) or [1-^14^C]oleate (C) and radioactivity incorporated in PC, PE and CL determined. Closed circles, control; Open circles, PVR. Data represent the means ± SEM of n=3, *p<0.05.

Since the increase in L_4_CL did not explain the mitochondrial dysfunction, we examined the levels of cardiac mitochondrial respiratory complexes. No alteration in the abundance of mitochondrial complexes was observed in PVR pups compared to controls (Suppl. Fig. 1B). LysoCL accumulation is known to cause mitochondrial bioenergetic dysfunction (24). We observed elevated trilinoleoyl-lysoCL (L_3_-lysoCL) species in hearts of PVR pups compared to controls (Fig. 5E). Thus, an elevation in linoleate containing L_3_-lysoCL species could be linked to the observed cardiac mitochondrial bioenergetic dysfunction in PVR rat pups.

The reason for the elevation in PE was also examined. Incorporation of radioactivity into PE from [1,3-^3^H]glycerol was unaltered in isolated cardiomyocytes from PVR rat pups indicating that *de novo* synthesis was unaltered (Fig. 6B). To further confirm this, synthesis of PE from serine and ethanolamine were examined. There was no alteration in synthesis of PE from [^3^H]ethanolamine nor synthesis of PE or phosphatidylserine from [^3^H]serine in cardiac myocytes from PVR rat pups (Suppl. Fig 3A,B). In contrast, the increase in PE was due to an increase in [1-^14^C]oleate incorporation into PE by 6 h of incubation indicating increased synthesis from oleate (Fig 6C).

## Discussion

In this study, we examined how altered cardiac CL and phospholipid metabolism and mitochondrial dysfunction are associated with the perinatal cardiac pathophysiology of PVR in perinatal rat pups. We show that PVR in perinatal rat pups result in cardiac hypertrophy and ventricular dysfunction, cardiac myocyte mitochondrial dysfunction with altered FA substrate utilization and elevated CL, L_3_-LysoCL and PE which may contribute, in part, to the cardiac mitochondrial dysfunction.

In this study, we have characterized for the first time the cardiac lipid alterations that occur in pups of the *in utero* hypoxia and indomethacin-induced fetal PVR rat model (13, 14). There are several reports of indomethacin induced pulmonary hypertension in human newborns mediated by premature constriction of the fetal ductus arteriosus (25–28). In addition, indomethacin-mediated closure of the ductus arteriosus in fetal rats was shown to result in early-onset right ventricular hypertrophy (29). Perinatal pulmonary hypertension in rats is known to permanently modify the pulmonary vasculature (30, 31). Consistent with this we observed that pulmonary arteries from our pups exhibited increased medial thickness in the larger pulmonary arteries with no changes in their external diameter. Associated with the pulmonary arterial modification was a cardiac hypertrophy characterized by elevation in right ventricle dimensions including RVWd and RVWs. In addition, left ventricle elevations in IVSd, IVSs, LVWd and LVSs, and reduction in EF and FS were observed. The elevation in right ventricular dimensions were consistent with that previously reported in this model of PVR (32).

Alteration in substrate utilization is a hallmark of cardiac dysfunction in HF (33). Studies in animal models and in human pulmonary arterial hypertension suggest that there is increased glycolysis and a metabolic shift from oxidative mitochondrial metabolism to the less energy efficient glycolytic metabolism (34, 35). Although basal OCR was unaltered in isolated cardiac myocytes of PVR pups when glucose was used as substrate, we observed increased glycolytic capacity and glycolytic reserve suggesting the potential for increase in glucose utilization for ATP synthesis. When palmitate was used as substrate an increase in FA oxidation was accompanied by an increase in ATP-insensitive respiration in isolated cardiac myocytes of PVR pups.

Pressure induced cardiac failure in rodents is known to result in increased PE levels and cardiac dysfunction (22). In addition, right ventricular pressure overload in adult rats induced by 12 weeks pulmonary arterial banding resulted in elevations in PE (34). We observed elevated cardiac PE levels, with no changes in PC, in hearts of perinatal PVR rat pups. The observed reduction in cardiac PC/PE ratio is consistent with that seen in pressure-induced heart failure (22). The increase in PE was not due to increased *de novo* synthesis or decarboxylation from phosphatidylserine since PE synthesis from radiolabeled glycerol, ethanolamine or serine was unaltered. In contrast, pulse-labeling with [1-^14^]oleate revealed increased synthesis of PE from oleate was responsible for the accumulation of PE in isolated cardiac myocytes from PVR rat pups.

A key question is whether alteration in CL levels actually contribute to the development of pediatric HF (36). A number of adult animal and human studies have indicated that reduced CL and L_4_CL accompany the development of HF (4, 37, 38). The reduction in cardiac CL in many early studies of adult HF in rodents can be attributed to the feeding of defined diets which may subject these animals to accelerated HF (39). The animals used in our study were harvested by caesarean section and were not subjected to maternal feeding. It was recently demonstrated that the relative percentage of L_4_-CL was preserved in pediatric human congenital single ventricle heart disease samples relative to biventricular controls (40). In addition, differences in CL content were not observed in induced pluripotent stem cell cardiac myocytes prepared from control and pediatric dilated cardiomyopathy with ataxia syndrome patients (41). Another study observed that the CL profiles in pediatric HF were unique from those in adults and the authors of this study hypothesized that end-stage pediatric heart failure adaptive mechanisms to preserve L_4_CL content may be intact to a greater degree than that seen in adult heart failure (42). Our results support the above hypothesis as we observed increased CL and L4CL levels in the hearts of caesarean section harvested perinatal PVR rat pups. Although *de novo* synthesis of CL from glycerol was unaltered in cardiac myocytes of perinatal PVR rat pups, an increase in [1-^14^C]linoleate incorporation into CL was observed which would explain the accumulation of CL and L_3_-lysoCL. The elevation in L_4_CL might contribute to the observed increase in FA oxidation in isolated cardiac myocytes from these animals. However, the apparent compensatory increase in FA oxidation in isolated cardiac myocytes from PVR rat pups was accompanied by an increased state 4 (oligomycin-inhibited) respiration indicative of an elevated proton leak which likely contributes to the mitochondrial dysfunction.

Accumulation of lysoCL is known to result in mitochondrial bioeneregtic dysfunction by compromising the stability of the protein-dense mitochondrial inner membrane leading to a decrease in optimal respiration (24). Although we observed no alteration in the abundance of individual mitochondrial complexes subunits, the significant accumulation of L_3_-lysoCL observed in the hearts of perinatal PVR rat pups might contribute to the mitochondrial bioenergetic dysfunction through a morphological disturbance of cristae membrane structure.

A limitation of the use of this hypoxia-induced model of PVR is whether the observed effects on metabolism may simply be due to the hypoxia itself. However, while hypoxia may cause a global change in cardiac homeostasis, the perinatal rats used in our study exhibited the cumulative effects of pressure, and the additional increase in cardiac weight was due to the effect of increased afterload. In addition, it is possible that the metabolic effects we observed may be more prominent in the right ventricle than the left ventricle. However, metabolism is also abnormal in the left ventricle in pulmonary arterial hypertension (43). In summary, our data show for the first time that a perturbed CL and PE metabolism is associated with and may contribute, in part, to the mitochondrial bioenergetic and cardiac functional defects observed in the heart of the hypoxia and indomethacin-induced gestational model of perinatal PVR in rats.

## Acknowledgements

The authors wish to thank Marilyne Vandel for technical assistance. LKC was the recipient of a CIHR/HSFC IMPACT Fellowship. GMH is a Canada Research Chair in Molecular Cardiolipin Metabolism. VWD is the Allen Rouse-Manitoba Medical Services Foundation Basic Scientist. This research was supported by an Environments, Genes and Chronic Disease Canadian Institutes for Health Research (CIHR) Team Grant #144626, the Heart and Stroke Foundation of Canada, the Natural Sciences and Engineering Research Council (NSERC), Children’s Hospital Research Institute of Manitoba (CHRIM) and the University of Manitoba Research Grants Program (URGP).

---

**Supplementary Figure 1.**
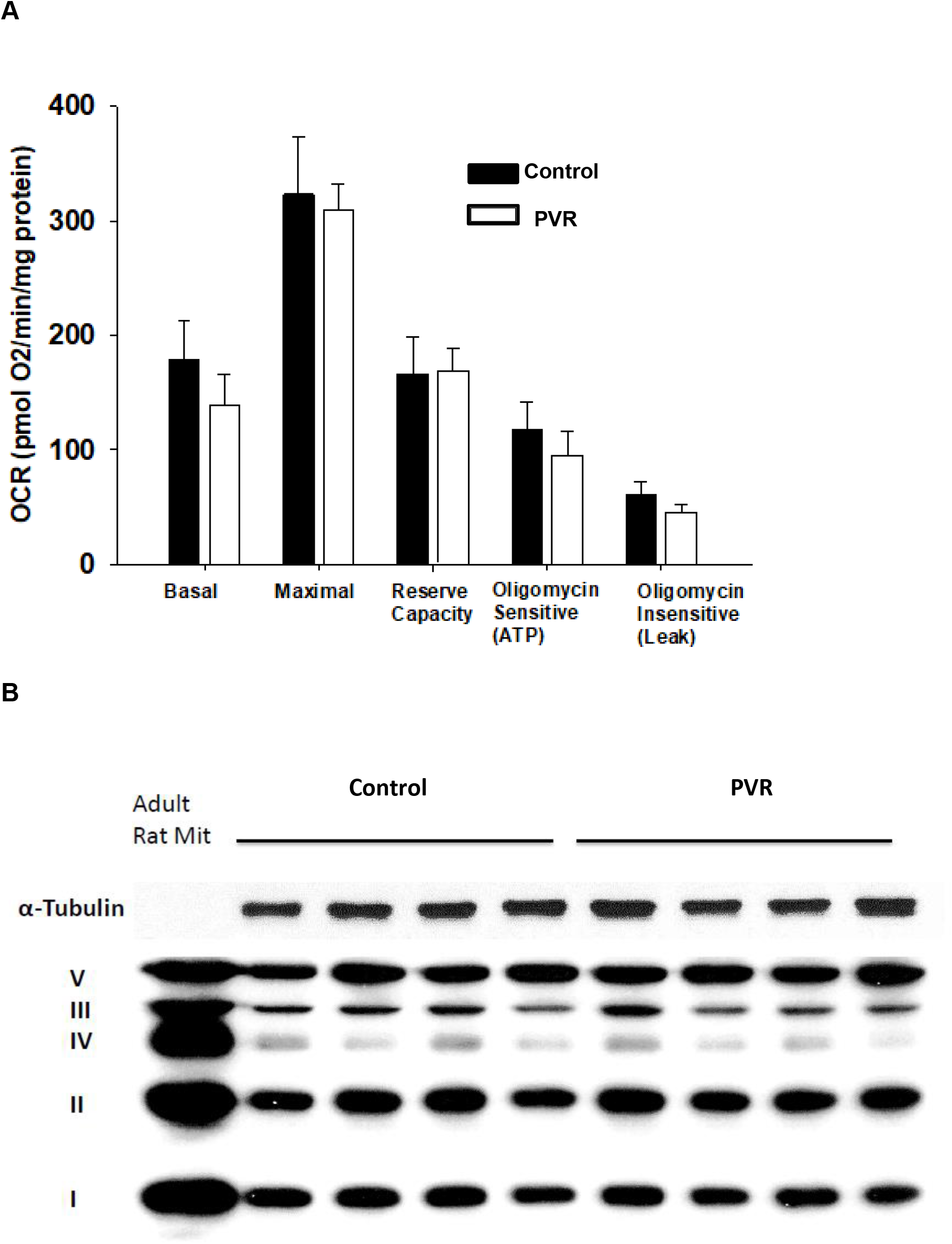
**A**. Mitochondrial oxygen consumption rate (OCR) in isolated control and PVR cardiac myocytes with glucose as substrate. **B**. Content of respiratory complexes in mitochondrial fractions from adult rat heart mitochondria and from control and PVR hearts. α-Tubulin is shown as loading control and individual respiratory complexes are indicated on the left. n=3-4.

**Supplementary Figure 2.**
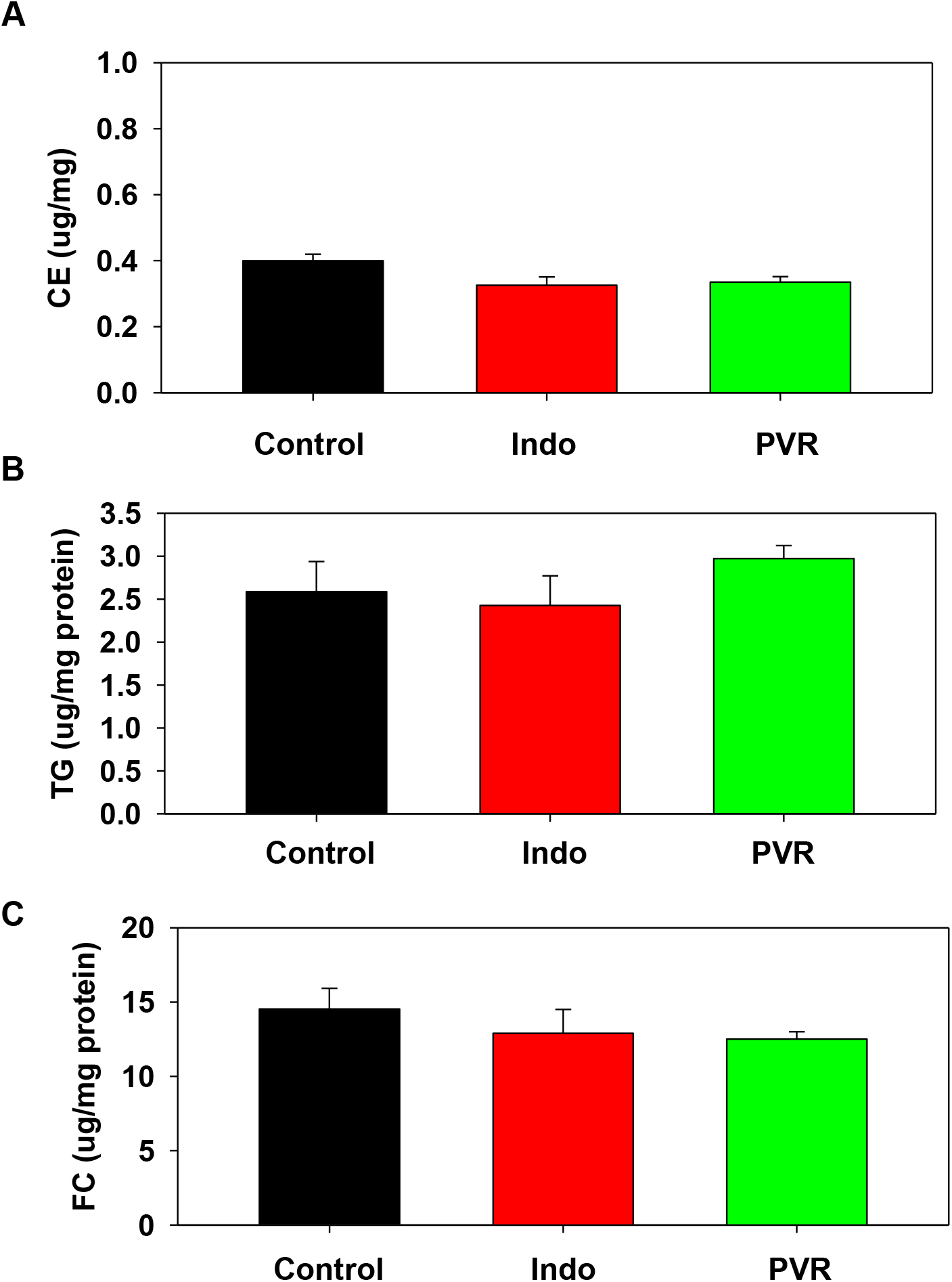
**A**. Cholesterol ester (CE); **B**. Triacylglycerol (TG); **C**. Free cholesterol (FC) in hearts of newborn pups from control, indomethacin (Indo) or PVR rats. n=3.

**Supplementary Figure 3.**
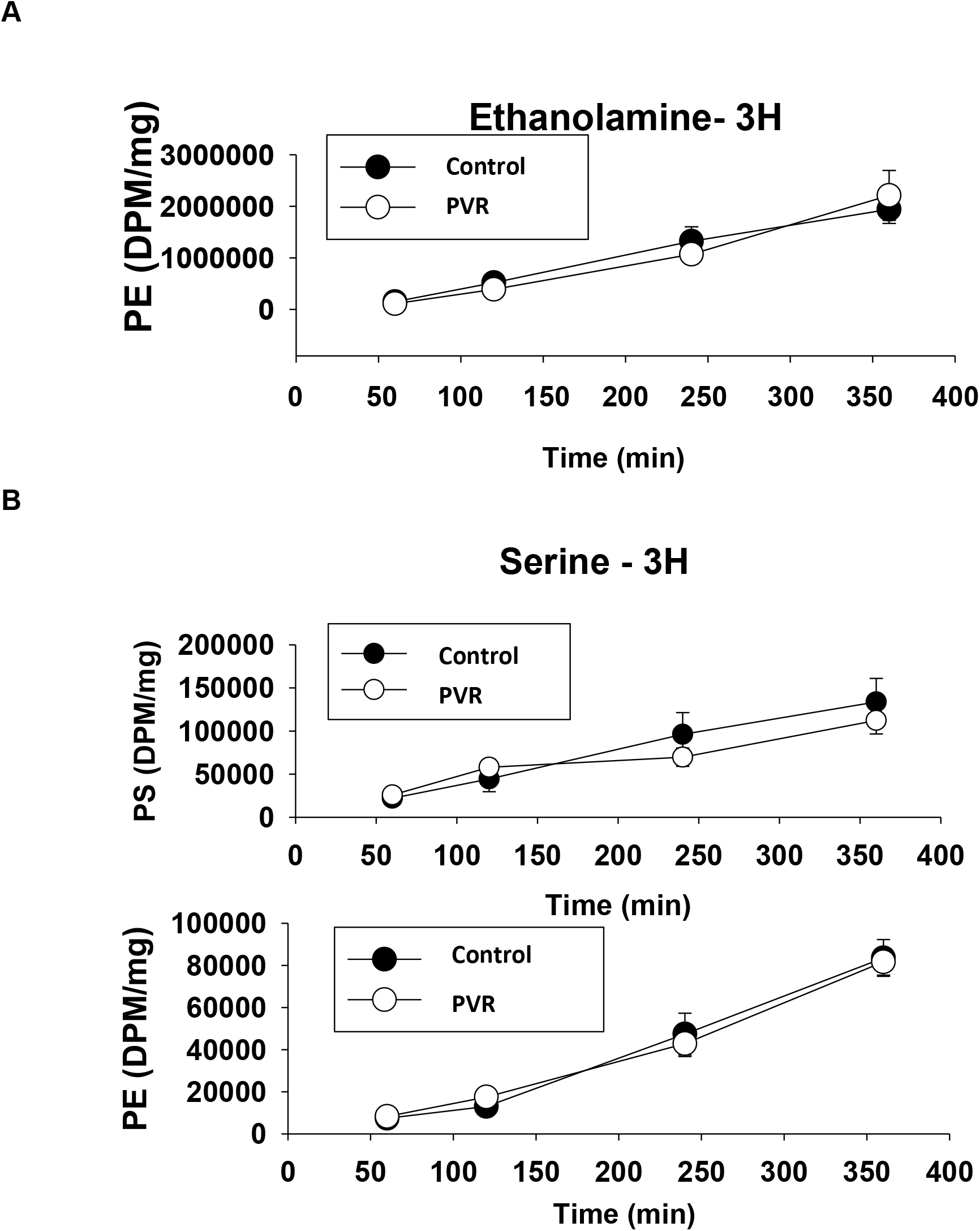
Synthesis of PE from ethanolamine or serine. **A**. Incorporation of [^3^H]ethanolamine into PE in isolated control and PVR cardiomyocytes. Incorporation of [^3^H]serine into phosphatidylserine (PS) or into PE in isolated control and PVR cardiomyocytes. n=3.

